# Identifying environmental factors affecting the production of pea aphid dispersal morphs in field populations

**DOI:** 10.1101/2023.03.23.534006

**Authors:** Michael J Bosch, Anthony R Ives

## Abstract

1. Many species of insects show phenotypically plastic dispersal traits in response to suboptimal environmental conditions. These polyphenetic traits can have large impacts on the population dynamics of the dispersing insects. Therefore, it is important to understand what environmental factors affect the development of dispersal traits and how these factors affect population dynamics via immigration/emigration.
2. Pea aphids (*Acyrthosiphon pisum*) exhibit a polyphenetic wing dimorphism in which individuals are either winged (alate) or not (apterous). The production of alate offspring is determined by environmental cues experienced by the mother. Environmental factors associated with these cues have been intensively studied, with crowding, host plant quality, natural enemies, and fungal infection being shown to affect the production of pea aphid alates under controlled laboratory conditions. Nonetheless, environmental factors affecting alate production have rarely been studied in field populations of pea aphids.
3. Using data from a three-year study of alate production in 5-9 alfalfa (lucerne) fields, we examined the effects of (i) pea aphid abundance, (ii) host plant maturity, (iii) temperature, (iv) predator and herbivore abundance, (v) parasitism, and (vi) entomopathic fungal incidence. In twice-weekly samples taken over the growing season, alate production ranged from 0 to 83% of the nymphal population.
4. Pea aphid abundance, temperature, and alfalfa maturity together explained 67% of the variation in alate production. The other factors we investigated explained little variation. These results suggest there is a limited number of key environmental factors that consistently predict changes in alate production in field populations, while many factors identified in lab studies may be unimportant. Our results highlight the value of investigating factors affecting the expression of plastic traits in insect species at a broad spatiotemporal scale and under natural conditions.

## Introduction

A diverse group of insect species exhibit environmentally determined morphotypes, or polyphenisms, for dispersal that take the form of flight-capable (typically winged) or flightless individuals (Braendle et al., 2006; Hochkirch & Damerau, 2009; Xu & Zhang, 2017). These dispersal polyphenisms are generally thought to be driven by trade-offs between potential reproduction and survival in temporally variable environmental conditions (Harrison, 1980; McPeek & Holt, 1992; Roff, 1986; Simpson et al., 2011; Zera & Denno, 1997). When environmental conditions are optimal in a patch of habitat, maximizing reproductive output is favored over dispersal if flightless individuals exhibit higher fecundity than winged individuals. As the condition of the patch becomes increasingly worse, dispersal is favored if it can lead to occupation of better habitat. A key question for dispersal polyphenisms is how individuals determine whether non-dispersing or dispersing morphs are produced. What environmental factors provide cues to individuals indicating when dispersal to a new patch will likely increase fitness?

Depending on how variable individual responses are, environmental cues can induce potentially large shifts in the phenotypic structure of a population, from flightless to flight-capable or vice versa (Crossley et al., 2022; Lagos-Kutz, 2020). These shifts can affect the population dynamics of the polyphenetic species by changing immigration or emigration between subpopulations in the surrounding area. A species with dispersal polyphenism could experience sporadic rates of gene flow, colonization of patches, disease transmission, and overall population growth depending on how spatially variable the environmental cues are (Hanski & Woiwod, 1993; Lago et al., 2020; Lin et al., 2018; Mazzi & Dorn, 2012; Simpson & Sword, 2008). Understanding what environmental factors are used for cues can help forecast the population dynamics of polyphenetic species and how the surrounding community will be affected (Sun et al., 2015; Wennersten & Forsman, 2012; Williams & Dixon, 2007).

The pea aphid (*Acyrthosiphon pisum* Harris) is a sap-feeding hemipteran that is an agricultural crop pest on several species in the Fabaceae family including peas, alfalfa, beans, and clover (Holman, 2009). During their parthenogenetic stage throughout the growing season, populations can double in size in as few as three days due to high fecundity and short development time (Harmon et al., 2009; Hutchison & Hogg, 1984, 1985). Pea aphids, along with most other aphid species, exhibit a polyphenetic wing dimorphism in their adult stage. The wing dimorphism is found only in parthenogenetic females during the summer growing season and is developmentally determined before birth by the mother depending on environmental cues she experiences (Müller et al., 2001; Sutherland, 1969a).

Environmental factors associated with changes in the propensity of pea aphid mothers to produce alates (winged offspring) has been intensively studied (Braendle et al., 2006). Crowding due to high pea aphid densities has been shown to increase alate production (proportion of offspring with wings) via an increase in tactile stimulation (antennal contact with other aphids) (Sutherland, 1969a). An increase in alate production in response to increasing host plant maturity has also been observed. This increase has been suggested to be a result either of increased movement leading to increased tactile stimulation or directly from the nutritional quality of the host plant (Sutherland, 1969b). Although the effects of temperature on alate production in pea aphids have not been directly investigated, studies conducted on other aphid species have shown an increase in alate production at lower temperatures. Much like the case for host plant quality, it is unclear whether the mechanism for alate response to temperature is direct or indirectly caused by increased tactile stimulation through changes in movement (Johnson, 1966; Schaefers & Judge, 1971). Natural enemies (predators and parasitoid wasps) have also been shown to increase pea aphid alate production, with natural enemy presence having a greater effect than factors associated with prior natural enemy presence (e.g., eggs, frass, and kairomones) (Dixon & Agarwala, 1999; Purandare et al., 2014). A wide range of natural enemy species has been investigated and results suggest that increased alate production can be attributed to the release of an aphid alarm pheromone ((E)-β-farnesene) causing an increase in aphid movement and inducing crowding effects (Hatano et al., 2010; Kunert et al., 2005; Podjasek et al., 2005; Sloggett & Weisser, 2002; Weisser et al., 1999). These effects reported for natural enemies are distinct from the ability of some parasitoid species to subvert wing development in parasitized juvenile aphids (Christiansen-Weniger & Hardie, 2000). Finally, pea aphid adults infected with fungal pathogens and colonies with fungus-infected individuals may have higher alate production due not only to increased movement and tactile stimulation but also to physiological cues expressed by infected individuals (Hatano et al., 2012). Although a diversity of factors can affect the propensity of pea aphid mothers to produce alates, crowding effects through tactile stimulation seem to play the largest role, with many other factors operating by changing the degree of tactile stimulation experienced.

Although alate production in pea aphids has been intensively studied, work has largely been confined to controlled conditions in laboratory settings where individual or colony responses to separate environmental factors are recorded. Little work has been conducted linking the results from these studies to alate production in the field, and therefore there is little evidence that cues affecting alate production in the lab have sufficient variation in the field to explain natural fluctuations in alate abundance. However, the wide range of factors shown to affect alate production creates a challenge in linking the results of lab studies to field observations, because many of the factors may be correlated in the field; for example, pea aphid abundance and natural enemy abundance are likely to be positively correlated, making it difficult to separate these two possible cues explaining variation in alate production. Nonetheless, the challenge presented by correlated explanatory variables can be addressed with appropriate statistical approaches.

In this study, we identified which of the cues that have been shown to affect pea aphid alate production in the lab can also explain variation in alate production in the field. Specifically, we analyzed the effects of pea aphid abundance, alfalfa height (as a measure of plant maturity), temperature, predators, parasitism, fungal infection, and herbivores on alate production in 5-9 alfalfa fields over three years. We included herbivore abundance even though no prior study had investigated their effects on alate production; this allowed us to compare herbivore abundance with predator abundance to see if competition or predation risk had differing effects. We used statistical methods designed to account for collinearity between predictor variables (environmental cues) and for spatiotemporal correlation between observations across the samples taken twice weekly from each field. Because previous studies showed or suggested crowding to be a major cue for alate production, we expected pea aphid abundance to be the strongest predictor of alate production in the field. This is the first study of which we know investigating the roles of multiple environmental factors on pea aphid alate production in field populations.

## Materials and methods

### Alfalfa field surveys

Pea aphid field surveys were conducted from 2017 to 2019 at the Arlington Agricultural Research Station (AARS) in Arlington, Wisconsin, USA, between 1 May and 1 September, typically twice per week (every 3-4 days). Surveys were conducted in 9, 6, and 5 fields in 2017, 2018 and 2019, respectively. Fields varied in the year they were initially planted and whether they were conventionally or organically managed. Harvesting occurred every 5-7 weeks, with surveys resuming in a field once the alfalfa reached approximately 12cm in height.

Survey samples in each field comprised six random transects, with insects collected along the transects using a 40cm diameter canvas sweep net. The number of sweeps per transect was determined by pea aphid abundance estimated with a single preliminary sweep, with the number of sweeps (5 - 150) selected to be roughly inversely proportional to pea aphid abundance. After sweeps along a transect were completed, pea aphids in the sweep net were counted along with Asian ladybeetle adults (*Harmonia axyridis*), seven-spotted ladybeetle adults (*Coccinella septempunctata*), spotted ladybeetle adults (*Coleomegilla maculata*), minute pirate bug adults (*Orius insidiosus*), nabids (*Nabidae spp*.), green lacewing larvae (*Chrysoperla rufilabris*), alfalfa weevil larvae (*Hypera postica*), alfalfa caterpillars (*Colias eurytheme*), potato leafhoppers (*Empoasca fabae*), and plant bugs (*Miridae spp*.). We used the number of insects per sweep as an index of abundance for pea aphids, predators, and other herbivores. Due to high variability in the abundance of the three ladybeetle species, we combined the three species to give a measure of overall adult ladybeetle abundance. To determine alfalfa regrowth following harvest, during each survey we measured the height of six randomly chosen alfalfa stems in each field from the base of the stem to the tip of the top leaflet.

### Parasitism and fungal infection samples

During field surveys, third and fourth instar pea aphids were aspirated from the sweep net samples and returned to the lab in a cooler to determine parasitism levels and fungal infection. The number of aphids per field sample was 15-100 depending on the aphid abundance. Within 48h of collection, aphids were dissected under a stereoscope, and the presence/absence of parasitoid wasp larvae and fungal mycelia were recorded. Counts of parasitized aphids were used rather than parasitoid wasp counts in sweep samples to provide a more accurate measure of the abundance of pea aphid parasitoids. An index of parasitism within a field was calculated as the number of parasitized aphids divided by the total aphids that were dissected. An index of fungal infection was calculated as the number of aphids with mycelia present divided by the total number dissected.

### Wing bud samples

We similarly sampled aphids for presence/absence of wing buds. Between 15 and 100 third and fourth instar were returned to the lab and within 48 hours were identified as having wing buds or not. The proportion of juvenile third and fourth instars with wing buds was used to measure alate production, rather than counts of winged adults, to exclude immigrant winged adults.

### Daily maximum temperature

Data on daily maximum temperature for AARS were collected from the University of Wisconsin Extension Ag Weather Program in 2017 and the Michigan Automated Weather Network and Enviro-weather Program in 2018 and 2019. Temperature data were collected from 1 May to 1 September for each year.

### Statistical analyses

To analyze the effects of pea aphid abundance, predator abundance, parasitism, fungal infection, herbivore abundance, temperature, and alfalfa height on alate production, we used a linear mixed model with autoregressive-moving average errors (LMM-ARMA) to regress wing bud proportion against the predictor variables. Illustrated for a single predictor variable, the structure of the model is

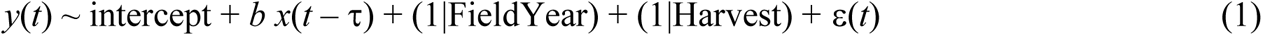

where *y*(*t*) is the arcsine-square-root-transformed proportion of aphids with wing buds in sample *t*, and *x*(*t* – τ) is the value of the predictor variable in sample *t* – τ where τ gives a time delay (number of samples) between the value of the predictor variable and when the proportion of wing buds in the population is likely to change in response. By standardizing values of *x*(*t* – τ) to have mean zero and standard deviation one, the regression coefficient *b* measures the effect size of predictor variable *x*(*t* – τ). To further investigate the conflicting effects of parasitoids on alate production (i.e., alarm pheromones versus subversion of wing development), a second analysis was conducted to factor out parasitized aphids that do not form wing buds due to physiological suppression by the parasitoid, the results of which are presented as Supplemental Information.

Because wing development depends on conditions experienced by the mother, we included the time delay τ between the proportion of alates and conditions experienced by their mothers. The effect of exposure on offspring wing development is greatest for offspring born 24 to 48 hours after maternal exposure, and the age of third and fourth instar pea aphids in field conditions ranges from 3 to 7 days, making the delay between exposure and sample collection between 4 and 9 days (Meisner et al., 2014; Sutherland, 1969a). Therefore, for the predictor variables temperature, alfalfa height, pea aphid abundance, predator abundance, herbivore abundance, and fungal infection, we used a delay of two samples (τ = 2) since two samples were taken per week. For parasitism rates determined from dissections, larval parasitoids were detected in third and fourth instars roughly 3-8 days after parasitoid attack, making parasitism samples representative of parasitoid abundance and activity 3-8 days prior (Henry et al., 2005; Meisner et al., 2014).

Due to the similar timing between parasitoid larval development and wing bud development, parasitism samples collected on the same day as wing bud samples were used in the analysis (τ = 0). Sampling was paused after harvest until sufficient alfalfa regrowth had occurred for sweep sampling. Therefore, wing bud samples in which there was no corresponding data for any of the predictor variables were removed from the analyses, resulting in 173 field samples in total.

Equation 1 contains random effects for field-years and harvesting cycles given by the variables FieldYear and Harvest, respectively. For the field-year random effect, each field-year was given its own value of FieldYear, and these values were assumed to have a Gaussian distribution with mean zero and variance estimated in the model fitting. To account for changes in the proportion of wing buds that coincide with harvesting events, each harvest cycle (the period between consecutive harvests) for each field-year was given its own value of Harvest which was assumed to have a Gaussian distribution with mean zero and variance estimated in the model fitting. Note that Harvest is a nested variable within FieldYear, because all samples with the same value of Harvest are contained within the same FieldYear.

The error term ε(*t*) in equation 1 accounts for both temporal and spatial autocorrelation in the data. To explain this term, we will modify the notation so that ε*i*(*t*) refers to the random error for field *i* in sample *t*. We assume that ε*i*(*t*) is given by a autoregressive-moving average process with autoregressive and moving average lags of 1 (i.e., an ARMA(1,1)). Thus,

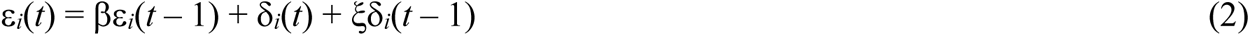

where β is the autoregressive parameter, and ξ is the moving average parameter. The Gaussian random variable δ*i*(*t*) is assumed to have no temporal autocorrelation, so that δ*i*(*t*) and δ*j*(*s*) for fields *i* and *j* (with *i* = *j* possible) are assumed to be independent for *t* ≠ *s*. Temporal autocorrelation occurs if a high value of ε*i*(*t*) in one sample is more likely to be followed by a high value in the following sample (β > 0) as might be the case if adult pea aphids that produced more winged offspring in sample *t* – 1 were more likely to continue producing more winged offspring in sample *t*. The moving-average process is included to account for possible effects of measurement error; measurement error can cause negative correlations between consecutive samples (ξ < 0). Spatial autocorrelation is incorporated by assuming that δ*i*(*t*) and δ*j*(*t*) for fields *i* and *j* are correlated, with the correlation depending on the distance between them. Specifically, we assume that the correlation between δ*i*(*t*) and δ*j*(*t*) equals (1 – *g*)exp(–*dij*/*r*) where *dij* is the geographical distance between fields *i* and *j* scaled so that *dij* = 1 for the most distant fields. The parameter *r* is the range that gives the rate at which the effect of distance on the correlation decreases; high values of *r* imply greater correlations between δ*i*(*t*) and δ*j*(*t*) at greater distances. The parameter *g* is the nugget that captures field-specific variation; the greater *g*, the greater the proportion of variation in δ*i*(*t*) that is local (i.e., uncorrelated with δ*j*(*t*), *j* ≠ *i*). Parameters *r* and *g* were estimated during the model fitting process. The LMM-ARMA model was coded as a special case of the function pglmm() in the R package ’phyr’ (Li et al., 2020).

Because pea aphid, herbivore, and predator abundances were most likely correlated, analyses were performed as two separate models. A full model was used that regressed wing bud proportion against the main effects of maximum daily temperature, alfalfa height, parasitism, fungal infection, herbivore abundances (potato leaf hopper, plant bug, alfalfa weevil, and alfalfa caterpillar), and predator abundances (minute pirate bug, lady beetle, lacewing, and nabid).

LASSO regression was used on the full model to identify variables having the most influence on wing bud proportion while simultaneously minimizing the effects of multicollinearity (Dormann et al., 2013; Zhang et al., 2020). LASSO regression introduces a penalty term λ to the likelihood function, making it possible to identify less influential predictor variables while highlighting more influential variables (Hastie et al., 2009). A second, reduced model containing only the more influential predictor variables (as determined from LASSO regression) and their interaction effects was used to investigate the effects of these variables on alate production. To investigate the variation in the data explained by each predictor variable for both the full and reduced models, the partial *R*^2^ values were calculated from Ives (2019) using the maximum likelihood estimates from models with and without the selected predictor variable:

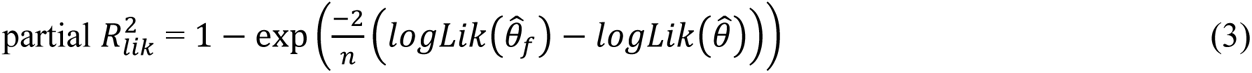

with *n* being the sample size of the dataset and *logLik*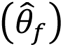 and *logLik*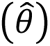 being the log-likelihood estimates for models with and without the predictor variable, respectively (Anderson-Sprecher, 1994; Edwards et al., 2008).

## Results

### Alfalfa survey data

To visualize the overall pattern of alate production and predictor variables through time, we averaged survey values among fields and across three consecutive sample days over the three years (Fig. 1). Alate production closely followed the population dynamics of pea aphid abundance across all three years (Fig. 1a). Harvesting events were largely synchronized among fields within the same year, with alate production often increasing during latter half of harvest cycles, but not for every cycle. Maximum daily temperatures fluctuated between 20°C and 30°C, with temperatures in 2017 and 2019 peaking in late-June to July; 2018 showed weaker seasonality. Alate production in 2017 and 2019 was highest in early and late summer, mirroring the pattern of temperature, although abundances followed a similar pattern making it hard to visually determine whether temperature, abundance, or both might affect alate production (Fig. 1b). Nabid and minute pirate bug abundances were the highest of the four predator groups, with peaks in abundances for all four groups tending to coincide with periods of increasing alate production (Fig. 1c, d). Plant bugs and potato leaf hoppers had the highest abundances of the four herbivore groups, with peaks in potato leaf hoppers occurring in July and peaks in abundances for the remaining three groups coinciding with periods of increasing alate production (Fig. 1c, d). Rates of parasitism often increased following peaks in alate production, while fungal infection varied greatly and only weakly coincided with increases in alate production (Fig. 1e).

**Fig. 1.**
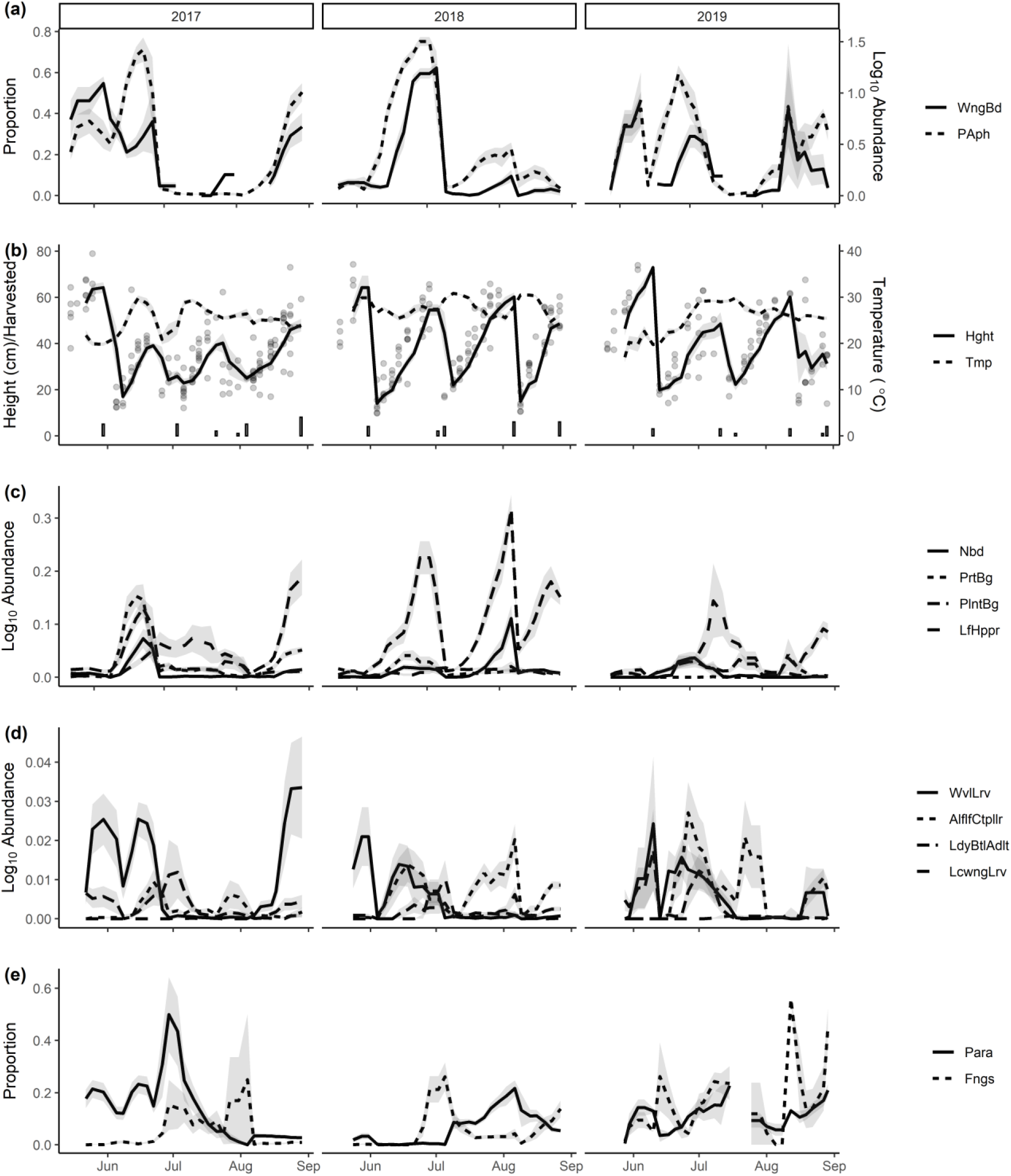
Field samples averaged from all fields for intervals of three consecutive sample days, with shaded areas giving ±1 SE. (**a**) Proportion winged pea aphids (WngBd) and log10 aphid abundance (PAph) averaged per sweep (with 5-100 sweeps taken per field sample). (**b**) Maximum daily temperature (°C) (Tmp) and alfalfa height (cm) (Hght), with points showing average height per surveyed field and bars along the horizontal axis marking the number of fields harvested following a given survey day. (**c**) Log10 abundances of nabid (Nbd), minute pirate bugs (PrtBg), plant bugs (PlntBg), and potato leafhoppers (LfHppr) averaged per sweep. (**d**) Log10 abundances of alfalfa weevil larvae (WvlLrv), alfalfa caterpillars (AlflfCtpllr), ladybeetle adults (LdyBtlAdlt), and green lacewing larvae (LcwngLrv) averaged per sweep. (**e**) Proportion parasitism (Para) and fungal infection (Fngs) as determined by dissections of 15-100 aphids per field sample.

### Full LMM-ARMA model with LASSO

Results from the full LMM-ARMA model showed a mix of positive and negative effects on the proportion of winged pea aphid juveniles, with pea aphid abundance and maximum daily temperature explaining a large amount of the variation (Fig. 2). Pea aphid abundance within fields had a strong positive effect on wing bud proportion (*b* = 0.25, partial 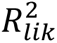 = 0.43), with weaker positive effects from alfalfa height (*b* = 0.06, partial 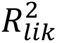 = 0.08) and abundances of plant bugs (*b* = 0.12, partial 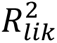 = 0.09), alfalfa caterpillars (*b* = 0.03, partial 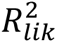 = 0.04), and ladybeetle adults (*b* = 0.05, partial 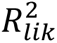 = 0.06). Negative effects occurred for maximum daily temperature (*b* = -0.07, partial 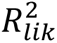 = 0.16), minute pirate bug abundance (*b* = -0.07, partial 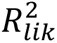 = 0.09), and parasitism (*b* = -0.06, partial 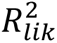 = 0.12). The effects of lacewing, leafhopper, alfalfa weevil, and nabid abundances were non-significant along with fungal infection. Results from the full LMM-ARMA model using the proportion of non-parasitized aphids with wing buds showed the same overall patterns, only with a reduced but still significant negative effect of parasitism (*b* = -0.04, partial 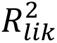 = 0.03; Fig. S1).

**Fig. 2.**
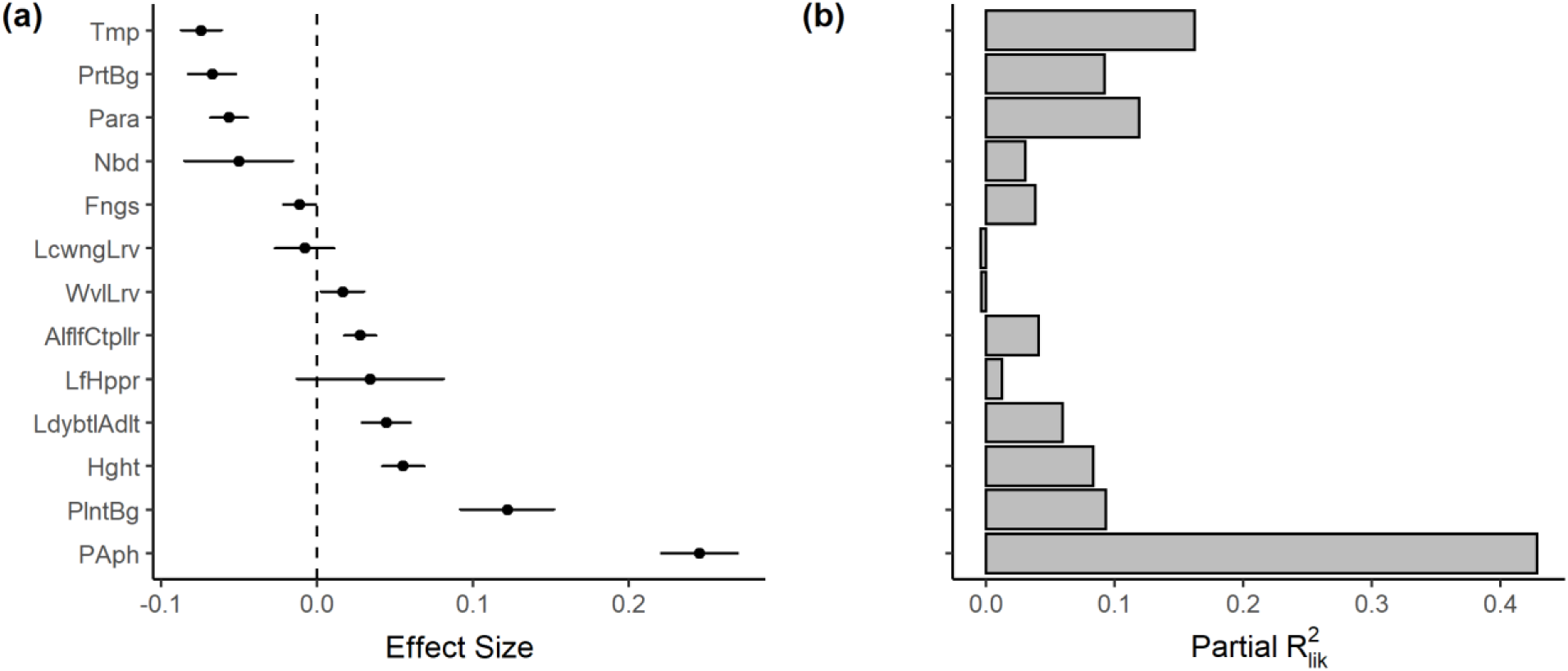
(**a**) Coefficient estimates (±1 SE) of the standardized effect size and (**b**) partial 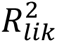 values for all predictor variables on the proportion of pea aphid juveniles with wing buds. Results are for the full LMM-ARMA model without LASSO. For abbreviations, see Fig. 1.

The correlations between predictor variables through time were sufficient to create a statistical challenge to identify the important factors causing variation in alate production; this is evidenced by the lack of correspondence between many of the variable effects sizes (Fig. 2a) and their corresponding partial 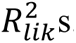. Except for pea aphid abundance and temperature, the partial 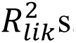 were low (Fig. 2b). The low partial 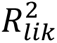 values imply that, when an independent variable is removed from the statistical model, other independent variables absorb the variation it explained. Thus, distinguishing the effects of the independent variables is difficult.

LASSO regression is designed to identify the most important predictor variables in a model when predictor variables are correlated (Dormann et al., 2013; Hastie et al., 2009; Zhang et al., 2020). Increasing the likelihood penalty λ decreased coefficient estimates of all predictor variables effectively to zero except pea aphid abundance, alfalfa height, and maximum daily temperature (Fig. 3), with a similar outcome using the proportion of non-parasitized aphids with wing buds (Fig. S2). The results from the LASSO regression were similar to the conclusions that were drawn from the partial 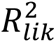 values, identifying pea aphid abundance and temperature as the most important independent variables (Fig. 2b). In addition, LASSO regression identified alfalfa height (maturity) as important, although less so than aphid abundance and temperature.

**Fig. 3.**
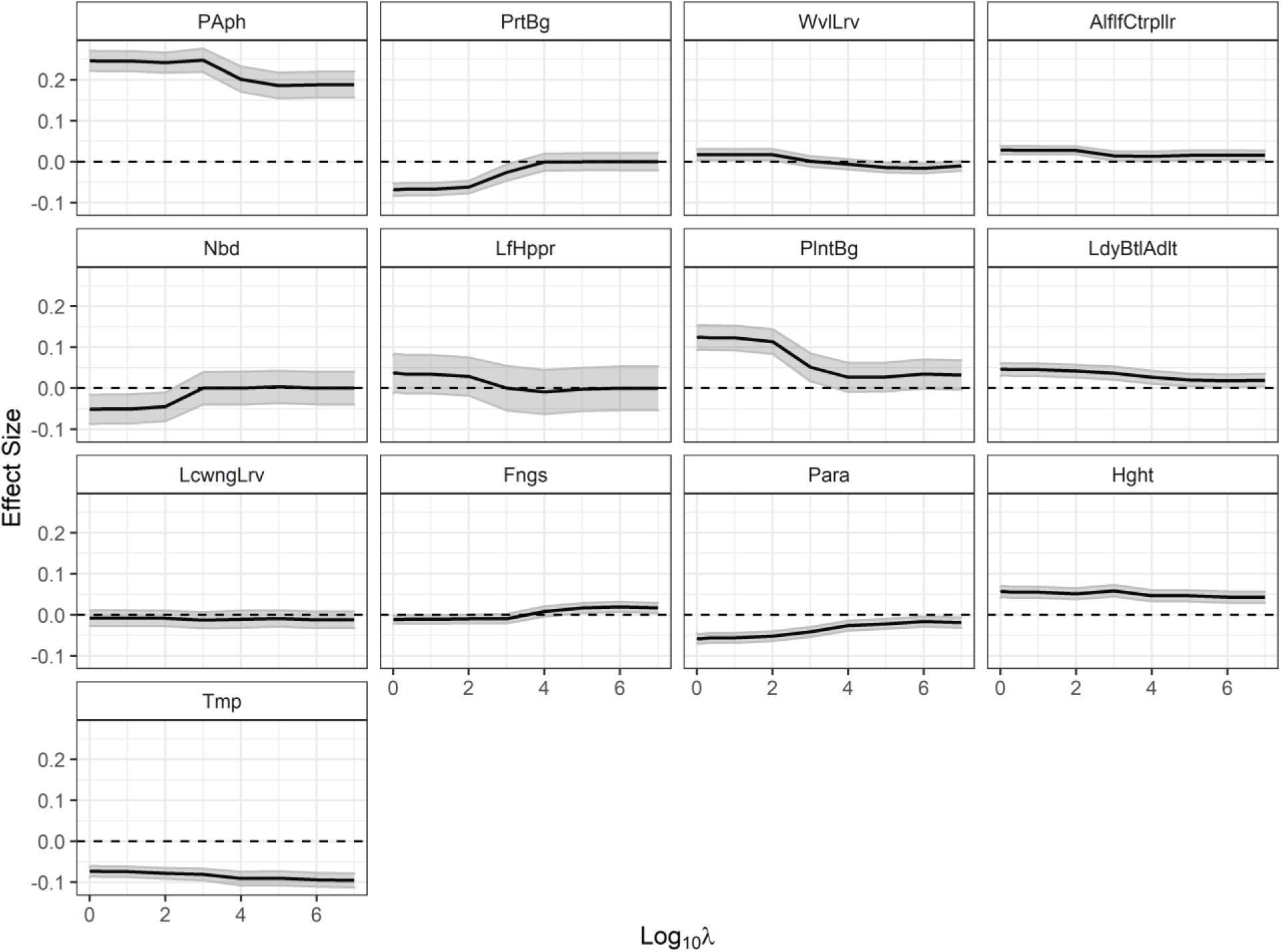
Coefficient estimates (±1 SE in gray) of all predictor variables in the full LMM-ARMA model using LASSO over log10 incremental increases in the penalty parameter λ. For abbreviations, see Fig. 1.

### Reduced LMM-ARMA model

The reduced LMM-ARMA model included those predictor variables identified in the LASSO regression: pea aphid abundance, alfalfa height, and maximum daily temperature. The reduced model also included all two-way interactions between predictor variables. Like the full LMM- ARMA model, pea aphid abundance and maximum daily temperature explained a large amount of variation in proportion wing buds (Fig. 4b). Both pea aphid abundance (*b* = 0.34, *P* < 0.0001, partial 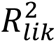 = 0.54) and alfalfa height (*b* = 0.05, *P* < 0.0001, partial 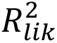 = 0.10) showed positive main effects on wing bud proportion, whereas maximum daily temperature showed a negative main effect (*b* = -0.12, *P* < 0.0001, partial 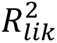 = 0.35). The interaction effects of maximum daily temperature on pea aphid abundance (*b* = -0.10, *P* < 0.0001, partial 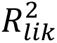 = 0.12) and alfalfa height (*b* = -0.07, *P* < 0.0001, partial 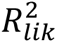 = 0.14) were both negative and similar in magnitude.

**Fig. 4.**
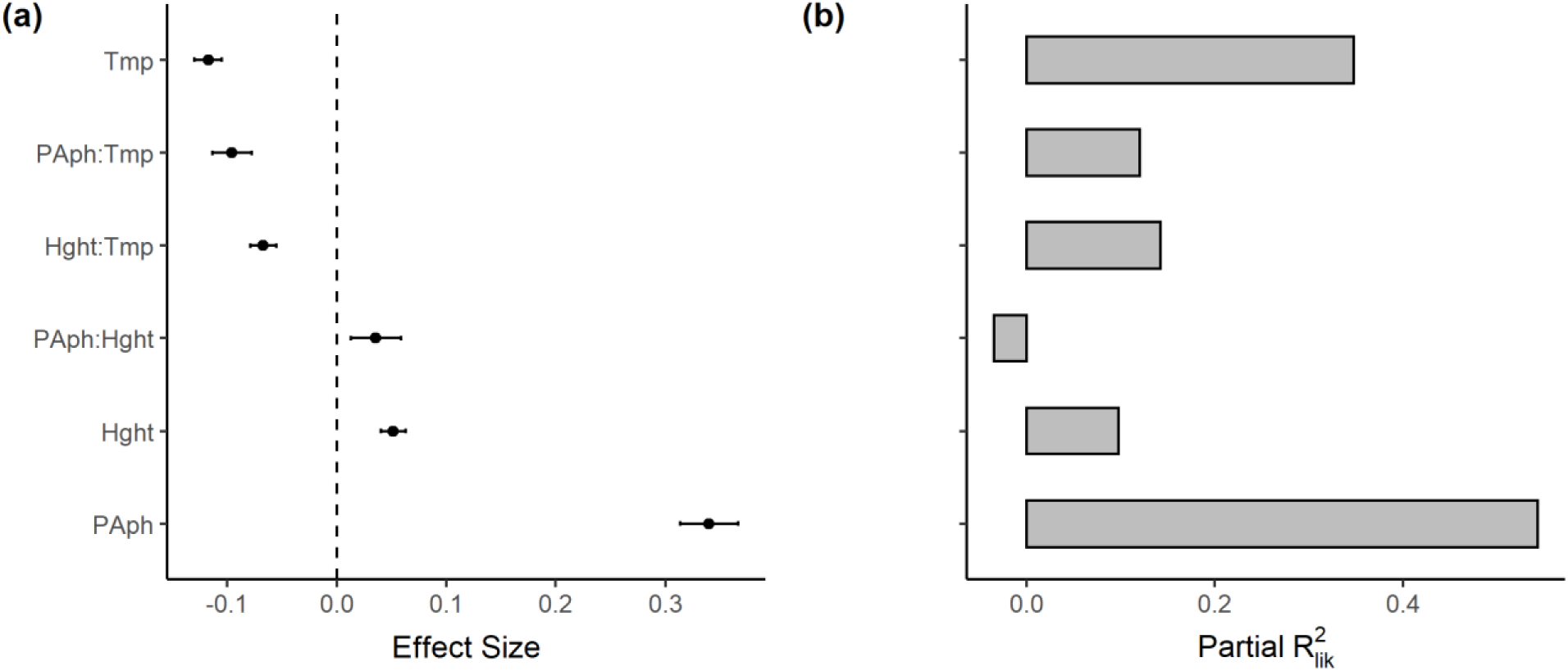
(**a**) Coefficient estimates (±1 SE) and (**b**) partial 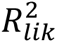 values for the main and two-way interaction effects of pea aphid abundance, alfalfa height, and maximum daily temperature on the proportion of pea aphid alates in the reduced LMM-ARMA model. For abbreviations, see Fig. 1.

The majority of partial 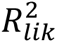 values in the reduced model were relatively high, implying that the reduced model explained a substantial portion of the variation in alate production.

Both full and reduced models showed moderate spatial effects in field-to-field variation; the nugget *g* was 0.79 and 0.76 in the full and reduced models, implying that 79% and 76% of the variance was "local" and not spatially autocorrelated (Table 1). Nonetheless, the remaining 21% and 24% of the variance was strongly correlated across fields, and the high values of the range parameter (*r* = 0.94 and 1.4 in full and reduced models) imply that spatial autocorrelation did not decrease much with distance between fields. These values of *g* and *r* imply that the correlations in residual errors for full and reduced models were 0.30 and 0.39 (equation used to calculate this is shown in Table 1). Temporal autocorrelation in both models given by the autoregressive (β) and moving average (ξ) parameters was low (Table 1).

## Discussion

The environmental factors associated with the production of alate offspring in pea aphids have been extensively studied under controlled lab conditions, with a wide array of different factors affecting the expression of this plastic trait (Braendle et al., 2006; Müller et al., 2001). Whether these results predict changes in alate production under natural conditions in field populations, however, depends on the relative degree of response to the different factors and the range of values of these factors that aphids experience under natural conditions. Here, we showed that abundance was the main predictor of alate production (proportion of juveniles with wing buds), which was consistent with findings under more controlled conditions (Purandare et al., 2014; Sutherland, 1969a). We also showed that alate production increased with the maturity of alfalfa, as has been found for pea aphids in previous studies (Sutherland, 1969b), and decreased at higher temperatures, which has been found for other aphid species (Johnson, 1966; Schaefers & Judge, 1971).

In contrast to previous lab experiments, however, we found no strong statistical support for association between alate production and the abundance of natural enemies, parasitism, fungal infection, or other herbivores. This might be explained because abundances of these groups were not large enough to elicit a strong aphid response. While changes in these factors may have corresponded with changes in alate production, these associations rarely or infrequently occurred over the span of this study (Fig. 1). Alternatively, this might be explained because we aggregated data on natural enemies, herbivores, and fungal mycelia at the field scale, which could mask any responses at the scale of single aphid colonies (Evans & Toler, 2007; Olson et al., 2000; Schooler et al., 1996). Thus, while natural enemies, parasitism, fungal infection, and other herbivores could potentially elicit a response in alate production, they do not explain a significant portion of the variation in alate production over multiple fields and years.

### Pea aphid abundance

Pea aphid abundance had the largest effect on alate production of the factors we investigated as measured by its partial 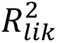 in both the full LMM-ARMA model that contained all predictor variables and the reduced model that contained only pea aphid abundance, alfalfa height, and temperature. Furthermore, in the LASSO regression, the coefficient for aphid abundance remained as the penalty parameter λ increased. Across all three years, alate production closely followed increases in pea aphid abundance (Fig. 1a). This pattern was not seasonal as there were two peak abundances in 2017 and 2019, but only one in 2018, yet alate production still followed aphid abundance. This pattern of alternating years with one and two peaks in pea aphid abundance was also documented by Meisner et. al. (2014) in the 1990s at this study site; they further showed with a model parameterized from lab population studies that the alternating one-peak/two-peak pattern is consistent with long-term host-parasitoid cycles in which there are three population cycles every two years. In our analyses, the effect of aphid abundance is present in the statistical model that includes parasitism, implying that the effect of fluctuations in aphid abundance are directly responsible for alate production rather than parasitism that may be responsible for the long-term cycles in aphid abundance.

Previous studies investigating the role of pea aphid density on alate production have suggested increases in tactile stimulation as the main mechanism producing crowding effects (Kunert et al., 2005; Purandare et al., 2014; Sloggett & Weisser, 2002; Sutherland, 1969a). Many of these studies have used experimental densities of pea aphids much higher than what would normally be seen in natural populations due to the non-colonial distribution of pea aphids among host plants. For example, in the study by Purandare et al. (2014), ten adult pea aphids were placed in a 32.5mm x 15mm Petri dish to increase tactile stimulation and induce alate production. Comparing this to the abundance of pea aphids in alfalfa fields collected from sweep nets and stem counts, catching one aphid per sweep corresponds to counting one aphid per approximately 150 alfalfa stems, or one aphid per 0.35m^2^ (Fig. S3). Therefore, pea aphid abundances that are likely to generate at least moderate alate production (0.5 log_10_ aphids per sweep; Fig. 1a) correspond to an average of only 0.014 aphids per stem. This makes our field results on the effects of pea aphid abundance difficult to explain in terms of tactile stimulation among aphids. Some clustering of pea aphids occurs among alfalfa stems; for example, first, second, and third instars are more likely to be found on stems with their mother (Duff & Mondor, 2012). However, unless the movement of juveniles away from their mother decreases as the overall aphid population abundance increases, this clustering of "family groups" cannot explain the increase in alate production with increasing population abundance as the clustering of family groups would be the same regardless of the population abundance.

An alternative explanation to tactile stimulation for our field results is that alate production is stimulated by the alarm pheromone released by pea aphids. This could potentially increase alate production at much lower pea aphid abundances than tactile stimulation. However, even though Kunert et al. (2005) showed that alarm pheromone concentration can increase alate production, these authors concluded that this effect is caused by tactile stimulation: alarm pheromones increase aphid movement which in turn increases tactile stimulation. A second alternative explanation is that alate production is stimulated by cues through the host plant, and higher aphid abundance increases the chances that an aphid will feed on alfalfa stems from plants that have previously been fed upon by other aphids. No prior studies have shown direct evidence for this explanation in pea aphids, but it is a potential mechanism by which pea aphids could perceive changes in population abundance. In addition, some aphid species increase the nutritional quality (amino acid concentration) of their host plant as more individuals feed on a single plant (Sandström et al., 2000). If this were the case for pea aphids, when population abundance increases individuals may show preference for alfalfa stems that either are currently or were previously fed upon, thereby increasing clustering and tactile stimulation. Although our field study confirms the result from prior experiments that increasing aphid abundance increases alate production, it also sheds doubt on whether the mechanism underlying this effect in the field is tactile stimulation among pea aphids.

### Alfalfa height

Alfalfa height was shown to have a positive effect on alate production, but there was no statistical interaction between alfalfa height and pea aphid abundance in the reduced LMM- ARMA model. These results are contrary to what has been suggested in previous studies that host plant quality only mediates alate production by inducing crowding effects (Sutherland, 1969b) and supports the hypothesis that host plant age has a direct effect on alate production. However, due to the low partial 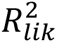 of alfalfa height in both the full and reduced model and the large negative interaction effect between alfalfa height and temperature, the direct effect of alfalfa height is small. Nevertheless, the effect of alfalfa height on alate production agrees with the nutritional quality of plant tissue typically decreasing as the host plant matures (Kennedy et al., 1950). For alfalfa specifically, protein content within a given stem decreases while lignin content increases with age, potentially resulting in amino acids becoming more difficult for pea aphids to obtain in older alfalfa (Fick & Mueller, 1989).

### Temperature

Maximum daily temperature had a strong negative effect on alate production with a high partial 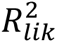, and there was a negative interaction between temperature and pea aphid abundance, implying that the negative effect of temperature was stronger at lower aphid abundance. These findings agree with what has been shown in other aphid species that experience higher alate production at lower temperatures (Johnson, 1966; Schaefers & Judge, 1971). Because temperature is seasonal, there is the possibility in the statistical models that temperature is acting as a surrogate for some other unmeasured variable that fluctuates seasonally. However, arguing against this are changes in alate production that were associated with non-seasonal changes in temperature. This can be seen during peak pea aphid abundances where prolonged lower temperatures in June of 2017 and 2019 were associated with greater alate production relative to pea aphid abundance than those seen in June of 2018, and August of 2017 and 2019 when temperatures were higher (Fig. 1a, b).

One explanation for the negative interaction between temperature and aphid abundance raises from the hypothesis that warmer temperatures increase the nutritional quality of plants, resulting in less clustering on stems that have been heavily fed upon by other aphids and thereby reducing crowding effects (Purwin et al., 2014; Rosinger et al., 1984). An alternative explanation could be that high pea aphid abundance (or poor plant quality) acts as a primary environmental cue indicating sub-optimal conditions and temperature as a secondary cue. Thus, when a primary cue (aphid abundance) becomes more prominent in an alfalfa field, the secondary cue (temperature) may become less important. This could help explain why there was a significant main effect of temperature after accounting for its interactions with pea aphid abundance and alfalfa height, which goes against what has been suggested in previous studies that temperature mainly affects alate production indirectly through crowding effects or plant nutrition (Braendle et al., 2006; Müller et al., 2001). Other studies have hypothesized that increased alate production directly in response to lower temperatures for aphid species may act as an effective bet-hedging method to maximize long-term fitness in more seasonal environments when environmental cues such as crowding and poor plant nutrition are not present (Liu, 1994).

Finally, temperature could increase drought stress in plants which could in turn affects alate production. Pea aphid population growth is reduced under drought-stress conditions (Forbes et al. 2005, Barton & Ives 2014). Therefore, if drought-reduced plant quality at higher temperatures were important, the expected relationship between temperature and alate production would be positive, rather than the observed negative relationship.

### Statistical analyses

Our statistical analyses faced two challenges. First, sampling was in close proximity in space (fields were 0.3-4.6km from each other) and in time, which may have generated both spatial and temporal autocorrelation in the residual random errors. We explicitly accounted for both types of autocorrelation by assuming that the random errors were given by an ARMA process, and that random fluctuations were correlated among fields depending on the distances separating them.

The statistical models detected little temporal autocorrelation, implying that the predictor variables could capture the changes in alate production from sample to sample without residual lagged effects in alate production. Spatial autocorrelation was greater, with the Pearson correlations in residual random errors between the two most distant fields (4.6km) estimated as 0.30 and 0.39 in the full and reduced LMM-ARMA models, respectively. This implies that there are environmental factors that create correlations in the production of alates among fields that are not included in the models’ predictor variables. Examples of factors that were not explicitly investigated in this study but could potentially affect the spatial autocorrelation between fields include precipitation and synchronized harvesting events across alfalfa fields. These factors could generate spatially correlated production in alates either through broad-scale changes in environmental cues (precipitation) or broad-scale disturbance (harvesting), which has not been as heavily investigated in the context of alate production by pea aphids. Indeed, there may be opportunities to discover additional factors that affect alate production across alfalfa fields.

A second challenge in the statistical analyses was investigating the effects of multiple predictor variables on alate production when many of these variables were correlated. This made it difficult to separate the individual effects of the predictor variables. In particular, some predictor variables, such as plant bugs, showed large effect sizes in the full LMM-ARMA model but nonetheless had low partial 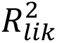. This means that even though the magnitudes of the effects of these predictor variables were large, removing them caused little loss in explanatory power of the model. Furthermore, the predictor variables that showed low partial 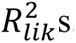 were also likely to be removed from the model in the LASSO regression. The advantage to using LASSO regression was that it removed predictor variables (such as minute pirate bugs and nabid) that had little individual explanatory power while retaining predictor variables like pea aphid abundance that have large individual effects on alate production.

## Conclusions

In this study, we have shown not only the main factors influencing alate production in field populations of pea aphids, but also the value of testing hypotheses from small-scale, manipulative laboratory experiments using field-scale observations under natural conditions (Levin, 1992). While previous work has shown a wide array of environmental factors that *can* affect alate production, our study has identified the environmental factors that *do* affect alate production in the field. The three factors pea aphids consistently respond to (diet, temperature, and pea aphid abundance) are factors commonly influencing dispersal polyphenisms in other insect species such as Roesel’s bush-cricket (*Metrioptera roeselii*) and the brown planthopper (*Nilaparvata lugens*) (Denno et al., 1986; Denno & Roderick, 1990; Hu et al., 2013; Sänger & Helfert, 1975; Uvarov, 1977; Zera & Denno, 1997). The frequency with which these three environmental factors appear to influence the expression of plastic dispersal traits in evolutionarily distinct organisms suggests that cues associated with diet, temperature, and density are reliable for maximizing individual fitness in the face of changing environmental conditions. Even though the specific mechanisms by which environmental factors affect the developmental switching in these different species may differ, the identity of the factors is the same. Identifying the key environmental factors eliciting polyphenetic dispersal traits is an important step in understanding the ecological and evolutionary consequences of dispersal for species in spatiotemporally varying environments.

## Acknowledgments

We are grateful to Ives lab members Salvatore DiVita, Sebastian Van Bastelaer, Megan Lipke, Lauren Godfrey, Ian Chen, Charlotte Dial, Leonardo Gargiulo, and Emily Gaare for the tremendous amount of field work they did, and to Jamie Botsch, Riley Book, Rachel Penczykowski, Lucas Nell, and Miriam Kishinevsky who over the years provided helpful feedback. This work was supported by NASA AIST program grant [80NSSC20K0282].

The authors declare no conflict of interest.

## Notes

https://doi.org/10.6084/m9.figshare.c.6463132

### Competing Interest Statement

The authors have declared no competing interest.

