## Supplemental Figures for "Identifying environmental factors affecting the production of pea aphid dispersal morphs in field populations"

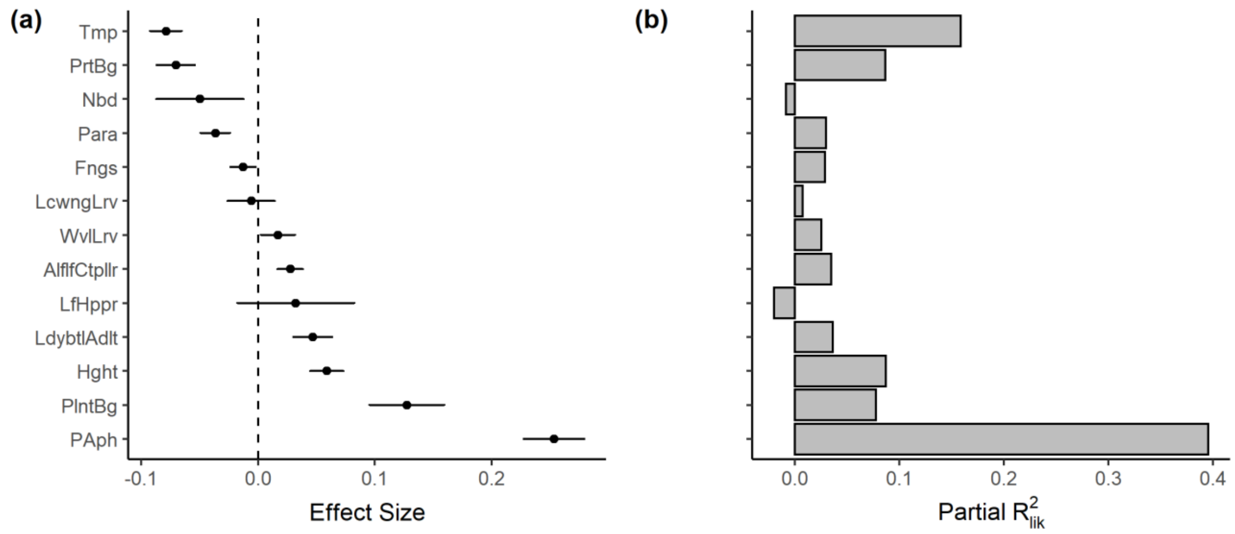


**Fig. S1.** (**a**) Coefficient estimates (±1 SE) of the standardized effect size and (**b**) partial $R^2_{lik}$ values for all predictor variables on the proportion of non-parasitized pea aphid juveniles with wing buds. This proportion was calculated by determining the proportion of aphids not parasitized within a wing bud sample (1 – index of parasitism) to derive the total number of non-parasitized aphids, and then taking the number of aphids with wing buds over this new total. Results are for the full LMM-ARMA model. For abbreviations, see Fig. 1.

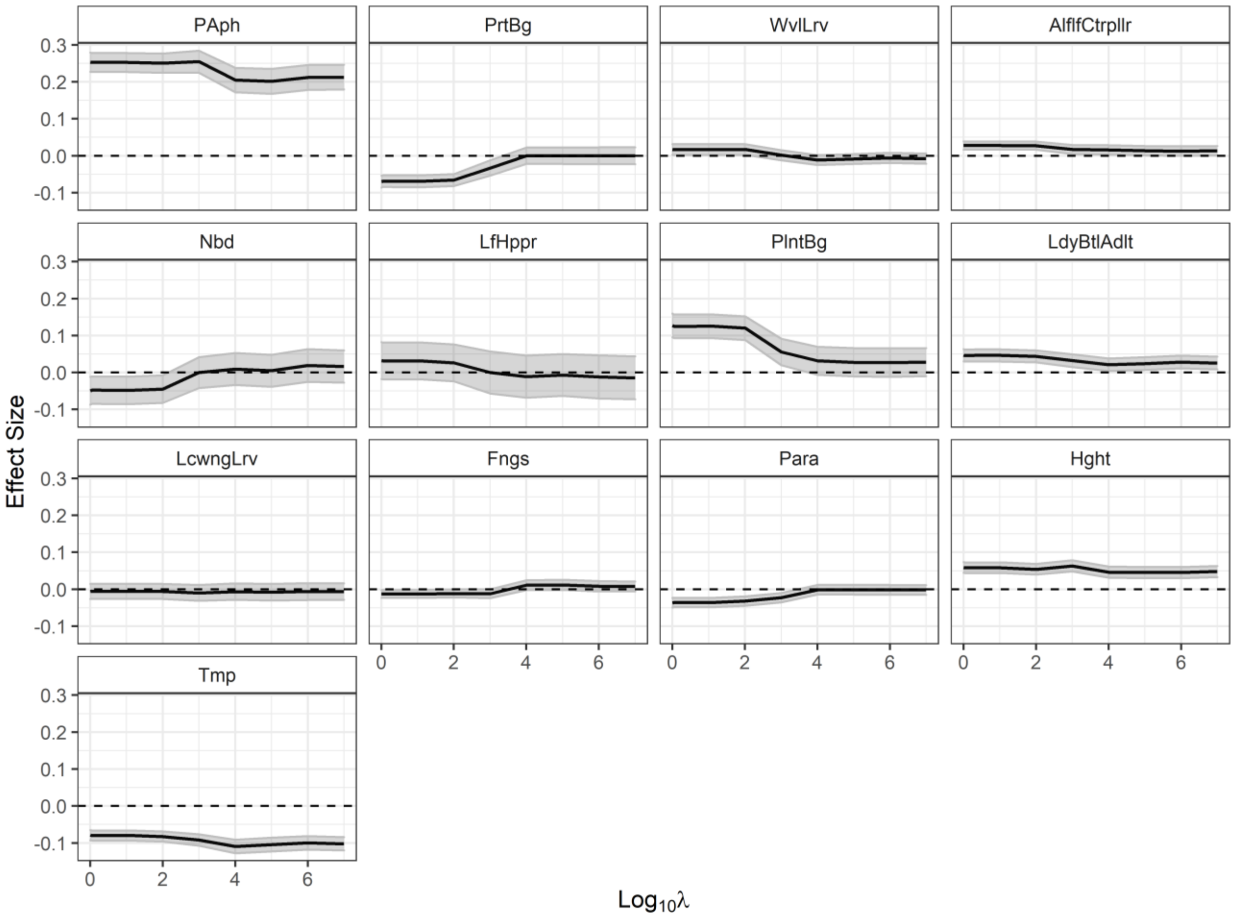


**Fig. S2.** Coefficient estimates (±1 SE in gray) of all predictor variables used on the proportion of non-parasitized pea aphid juveniles with wing buds in the full LMM-ARMA model using LASSO over $\log_{10}$ incremental increases in the penalty parameter $\lambda$. For abbreviations, see Fig. 1.

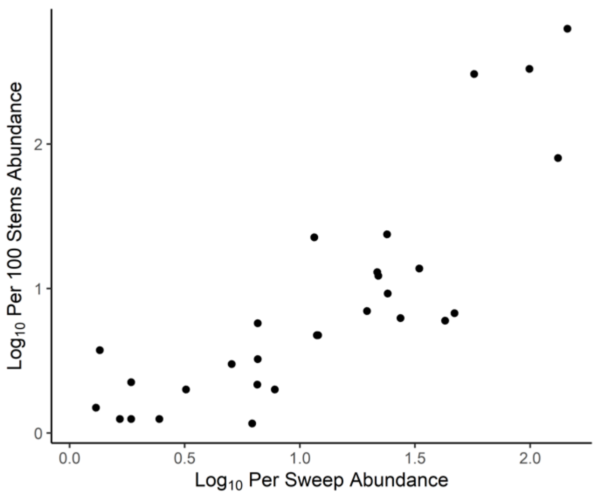


**Fig. S3.** Relationship between per sweep pea aphid abundance and pea aphid abundance per 100 stem counts within alfalfa fields. Sweep and stem count data used was collected during the summer of 1994.
